# microRNA profiling of mouse cortical progenitors and neurons reveals miR-486-5p as a novel regulator of neurogenesis

**DOI:** 10.1101/2019.12.27.889170

**Authors:** Martina Dori, Daniel Cavalli, Mathias Lesche, Simone Massalini, Leila Haj Abdullah Alieh, Beatriz Cardoso de Toledo, Sharof Khudayberdiev, Gerhard Schratt, Andreas Dahl, Federico Calegari

**Affiliations:** CRTD – Center for Regenerative Therapies Dresden, School of Medicine, TU Dresden; Fetcherstrasse 105, 01307, Dresden, Germany; DRESDEN-concept Genome Center c/o Center for Molecular and Cellular Bioengineering (CMCB), Technische Universität Dresden; Fetcherstrasse 105, 01307, Dresden, Germany; Institute for Physiological Chemistry, Biochemical-Pharmacological Center Marburg, Philipps-University of Marburg, Marburg, Germany

## Abstract

MicroRNAs (miRNAs) are short (∼22 nt) single-stranded non-coding RNAs that regulate gene expression at the post-transcriptional level. Over the past years, many studies have extensively characterized the involvement of miRNA-mediated regulation in neurogenesis and brain development. However, a comprehensive catalog of cortical miRNAs cell-specifically expressed in progenitor types of the developing mammalian cortex is still missing. Overcoming this limitation, here we exploited a double reporter mouse line previously validated by our group to allow the identification of the transcriptional signature to neurogenic commitment and provide the field with the complete atlas of miRNAs expression in proliferating neural stem cells, neurogenic progenitors and newborn neurons during corticogenesis. By extending the currently known list of miRNAs expressed in the mouse brain by over two fold, our study highlights the power of cell type-specific analyses for the detection of transcripts that would otherwise be diluted out when studying bulk tissues. We further exploited our data by predicting putative novel miRNAs and validated the power of our approach by providing novel evidence for the involvement of miR-486 as a novel player in brain development.

## INTRODUCTION

MicroRNAs (miRNAs) are short (∼22 nt) single-stranded non-coding RNAs that regulate gene expression at the post-transcriptional level (1,2). Canonical miRNAs derive from longer primary transcripts harboring a stem-loop that is processed by two RNAse III enzymes: Drosha in the nucleus and Dicer in the cytoplasm (3,4). Eventually, mature miRNAs are loaded into the RNA-induced silencing complex (RISC) (5,6) to destabilize or cleave complementary target messenger RNAs (mRNAs) thereby inhibiting their translation (7).

miRNA-mediated regulation of translation is far more than an adjustment of cellular protein levels, but rather an essential developmental mechanism. In fact, a number of mouse lines mutant for miRNA-processing enzymes or individual miRNAs showed dramatic phenotypes, ranging from impaired organogenesis to pre- and perinatal lethality (8-11). The effects of interfering with miRNA function were found to be particularly severe during brain development and leading to a decreased survival of neural progenitors and newborn neurons and ultimately causing cortical malformations (12-15). In addition, well-established regulatory loops are mediated by miRNAs such as in the synergistic effect of miR-9 and *let-7b* inducing neural progenitors differentiation by targeting the Tlx receptor (Nr2e1) (16,17) as well as down-regulating Hes1 and CyclinD1 as critical gene hubs controlling cell-cycle exit and enhancing differentiation (18,19). Moreover, it is well characterized the interaction of miR-9 with miR-124 to target the RE-1 Silencing Transcript factor (REST), a strong inhibitor of pro-neural genes (20-22). Many more examples are known of miRNAs controlling neurogenesis and brain development (23,24) highlighting the importance of studying their physiological expression patterns in different cell types of the developing cortex as a crucial step to gain insights into the pathways underlying their timely regulation and function. Remarkably, however, a comprehensive catalog of cortical miRNAs cell-specifically expressed in progenitor types and neurons is still missing.

The lack of a comprehensive catalog of miRNAs expression in specific populations of neural progenitor cells is due to many factors including technical limitation in the coverage of single-cell small RNA sequencing (25) and that essentially all previous high-throughput miRNA studies on neurogenesis used either microarrays or total brain lysates (26-30). As a consequence, the resolution of previous studies was limited by the variety of probes printed on the microarrays or, alternatively, by the coexistence in time and space of different cell types of the developing brain. To overcome these limitations, we here exploited a previously described dual-reporter mouse line, which allows the isolation of different neural progenitor types and newborn neurons (31).

More specifically, with the progression of neurogenesis two distinct, lineage-related populations of neural progenitors coexist in the developing cortex: radial glia, proliferative progenitors (PP) that expand the stem cell pool by symmetric divisions and neurogenic, differentiative progenitors (DP) that divide to generate neurons (N) (32,33). In studying the fate and nature of each population, several studies have identified the expression of defined molecular markers in each cell type. In particular, and by taking advantage of *Btg2* and *Tubb3* expression, our group has generated a combinatorial, double-reporter mouse line in which RFP and or GFP expression allowed the isolation specifically of PP, DP and N based on their endogenous fluorescence (RFP–, RFP+ and GFP+, respectively) (31).

Validation and use of this mouse line revealed to be very powerful in the identification of several new genes and biological processes regulating cortical development (31). This included the thorough characterization of the elusive class of long non-coding (34) and circular (35) RNAs, novel transcription factors involved in corticogenesis (36) and a comprehensive description of DNA methylation and hydroxymethylation as epigenetic marks tuning brain development (37).

Given the previously validated power of our approach, we here exploited this Btg2::RFP/Tubb3::GFP line to provide the field with a complete atlas of miRNAs expression in cortical progenitors and neurons of the mouse brain at embryonic day (E) 14.5 as a mid-stage of corticogenesis. Furthermore, and validating our approach, we provide evidence for the involvement of miR-486 as a novel regulator of corticogenesis.

## MATERIALS AND METHODS

### Animals and embryos dissection

Mice were housed into the Biomedical Services Facility (BMS) of the MPI-CBG under standard conditions (12-hour light-dark cycle, 22 ± 2°C temperature, 55 ± 10 % humidity, food and water supplied *ab libitum*). All experimental procedures were performed according to local regulations and all animal experiments were approved by local authorities (Landesdirektion Sachsen; 24D-9168.11-1/41, 2008-16, 2011-11, TVV 39/2015, 13/2016 TVV and 16-2018). Btg2^RFP^/Tubb3^GFP^ males were time-mated with C57BL/6J females, which were marked as E0.5 the morning that a spermatic plug was observed. Pregnant females were anesthetized using Isoflurane (Baxter) and sacrificed by cervical dislocation at E14.5. Brains of RFP/GFP double-positive embryos were collected and lateral cortices isolated after removal of meninges and ganglionic eminences. Plugged C57BL/6J females for *in utero* electroporation or RNA extraction for Northern blot were purchased from Janvier Labs. Mice were sacrificed at E14.5 or E15.5 and embryo brains and cortices were dissected as above.

### Cell dissociation and FAC-sorting

Lateral cortices of RFP/GFP double-positive embryos were dissociated using Papain-based Neural Tissue Dissociation Kit (Miltenyi Biotech) according to the manufacturer’s protocol. Cells were resuspended in 1 ml of ice-cold PBS and 10 µl of 7-AAD (BD Pharmingen) were added for dead cells discrimination. Sorting was performed by BD FACSAria™ III (BD Biosciences) with previously described gating (31,35). A minimum of 1 × 10^6^ cells per sample was collected in PBS and centrifuged (300 g, 10 min at 4°C) before RNA extraction.

### RNA extraction

For miRNA deep-sequencing, total RNA was isolated using Quick RNA Mini Prep kit (Zymo Research) from cells sorted as described above. RNA quality and integrity were assessed by Bioanalyzer (Agilent Genomics). RNA integrity values (RIN) were above 9.0. For Northern blots, total RNA was isolated by TRI Reagent (Sigma-Aldrich). Briefly, lateral cortices of all E14.5 embryos of one litter were pooled and lysed in 1 ml of TRI Reagent. Samples were added 200 µl of chloroform, mixed and left at RT for 15 min before centrifugation at 12,000 g for 30 min at 4°C. Aqueous phases were transferred to new tubes and RNAs were precipitated by adding 500 µl of 2-propanol. RNA pellets were washed with 1 ml of 75 % ethanol and eventually resuspended in 50 µl of nuclease-free water.

### Library preparation and small RNA deep sequencing

Library preparation was performed on 1 µg of total RNA with NEB Next Small RNA Library Prep Kit. All cDNA libraries were prepared according to the manufacturer’s specifications, including adapter ligation, first-strand cDNA synthesis, PCR enrichment and size selection. cDNA purity and concentration after gel extraction were measured by qPCR. Samples were sequenced on Illumina HISeq 2500 and single-end 75-bp reads were obtained.

### Bioinformatics and statistical analyses

Sequencing data were obtained for PP, DP and N in 3 biological replicates. After adapter removal, reads shorter than 30 bp were aligned to miRBase v.20 (38) using gsnap (39). Alignment was performed in 3 consecutive steps: a) on mature miRNAs sequences, b) unmapped reads were extracted and c) aligned on precursor-miRNA. During all steps, no mismatches were allowed and multi-mapped reads discarded. Eventually, a table of read counts per mature miRNA (read count ≥ 1) was assembled. For novel miRNA prediction, all unmapped reads were extracted and aligned using miRDeep2 (40) on mouse genome (mm10). The R package DESeq2 (41) was used for normalization of the read count table and further testing of differential expression. Mean counts from replicates were used for fold change (FC) calculations: log2FC values >= 0.58 or <= −0.58 were considered up- or down-regulation, respectively. Benjamini–Hochberg procedure was applied for multiple t-test adjustment and FDR values lower than 0.05 were considered significant. A minimum of 3 biological replicates was used for any other assessment presented in the manuscript. Statistical differences of mean values were calculated by two-tailed student t-test, assuming p<0.05 as significant.

### In utero electroporation

LNA oligonucteotides (miRCURY LNA miRNA Inhibitors) were purchased from Exiqon and co-electroporated with pDSV-mRFPnls reporter plasmid (Lange et al., 2009). LNA sequences are reported in **Supplementary Table 1**. *In utero* electroporation was performed as previously described (42,43): C57BL/6J pregnant mice were anesthetized with isoflurane at E 13.5 and 1 µl of DNA solution (10 µM LNA, 0.8 µM RFP plasmid) was injected into the embryo left ventricle, followed by the application of 6 electric pulses (30V and 50 ms each at 1 s intervals) through platinum electrodes using a BTX-830 electroporator (Genetronics).

### Immunohistochemistry

After dissection, brains were fixed in 4 % paraformaldehyde in 0.1M phosphate buffer (PFA, pH=7.4) overnight at 4°C, cryoprotected in 30 % sucrose and cryosectioned (10 µm thick slices). Immunohistochemistry was performed as previously described (42) (**Supplementary Materials and Methods** and **Supplementary Table 2** for a list of used antibodies). Nuclei were counterstained with DAPI. Sections were imaged using an automated microscope (ApoTome; Carl Zeiss), pictures digitally assembled using Axiovision software (Carl Zeiss) and composites analyzed using Photoshop CS6 (Adobe).

## RESULTS

### The comprehensive miRNome of neurogenic commitment

Aiming to profile global miRNA expression during cortical development, we isolated PP, DP and N (each in three biological replicates) from the lateral cortices of Btg2::RFP/Tubb3::GFP mouse embryos at E14.5, as previously described (31,35) (**Figure 1a**). Total RNA was used for cDNA library preparation and small RNAs were isolated by size selection, followed by 75-bp high-throughput sequencing. To assemble the catalog of cortical miRNAs, we aligned reads with gsnap (39) and used miRBase (v.20) as the most complete reference available to date (38) yielding an average of 1.5 million unique-mapped reads (51% of total). Within the mapped reads, we detected (defined to as reads ≥ 1) 1,058 mature miRNAs derived from 703 precursor transcripts (pre-miRNA) corresponding to 55% and 59% of the 1,908 mature and 1,186 pre-miRNAs reported in the reference, respectively. More specifically, 640 mature miRNAs were in common to all 3 cell-types while 49 (4.6%), 58 (5.5%) and 129 (12.2%) were specific to PP, DP and N, respectively (**Figure 1a**). Notably, when compared to a previous study which reported 294 pre-miRNAs (read count ≥1) expressed in the whole E15.5 mouse brain (30), our dataset included essentially all (96%) of these previously known cortical miRNAs and further doubled this list by including additional 421 pre-miRNAs. In turn, this highlights the power of cell type-specific analyses for the detection of transcripts that would otherwise be diluted out when studying bulk tissues.

**Figure 1.**
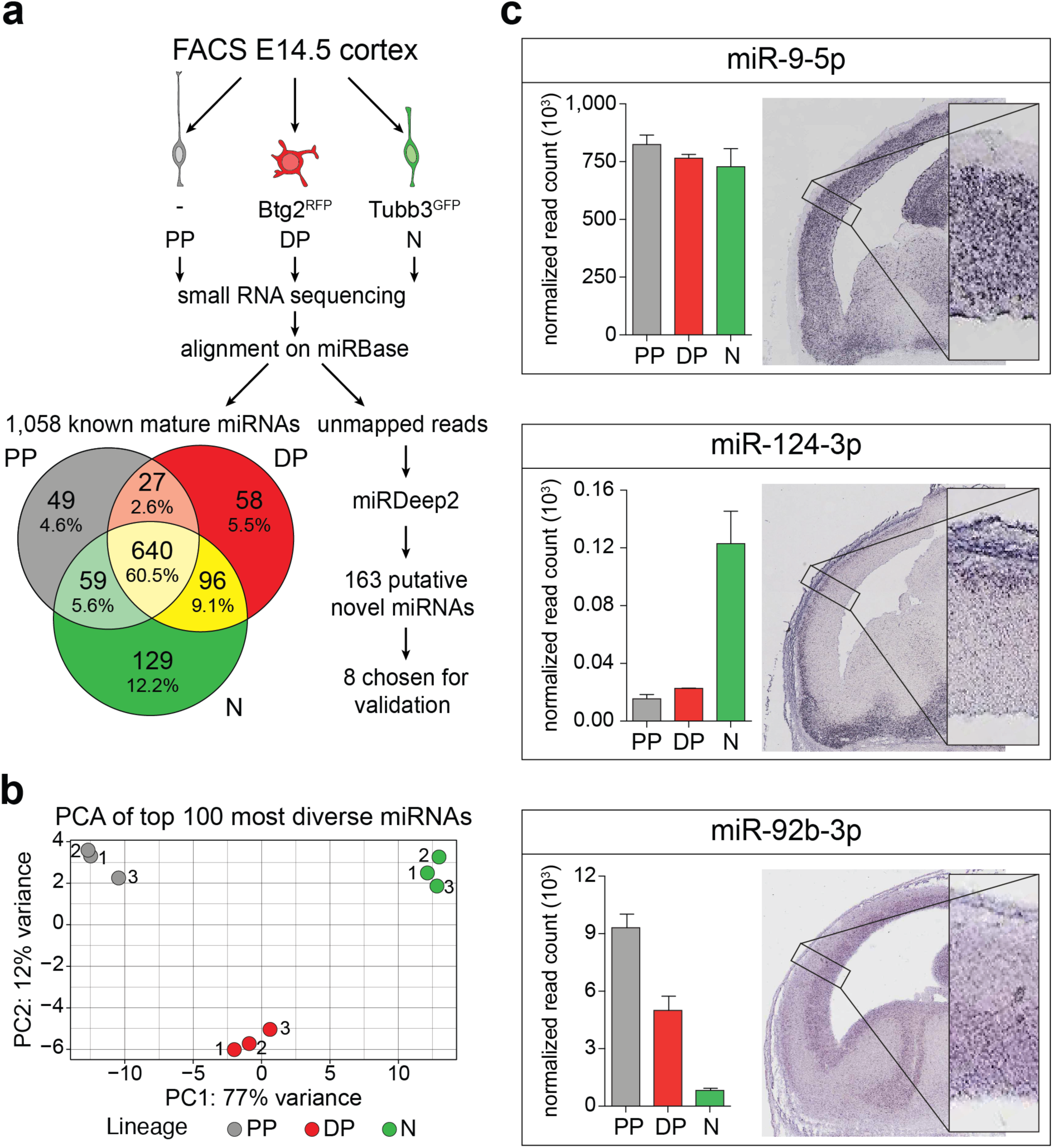
Assembly and validation of cortical miRNome. **a)** Outline of the steps taken to generate cortical miRNome: sorting of E14.5 PP, DP and N, followed by small RNA sequencing. Mature miRNAs were identified through alignment on miRBase and novel miRNAs were predicted by miRDeep2. **b)** Principal component analysis of DESeq2-normalized 100 most diverse miRNAs between biological replicates (1-3) and cell populations (proliferative progenitors, grey; differentiative progenitors, red; neurons, green). **c)** Sagittal sections of E14.5 cortices downloaded from Eurexpress. Magnifications of the lateral cortex are shown (bottom-right) to appreciate the extent of the overlap with miRNA expression data measured by deep sequencing (histograms). Error bars = s.d. N = 3.

Furthermore, given that 42% of our reads did not align to any known miRNA and recent studies reported the detection of novel miRNAs in both mice and humans (44,45), we hypothesized that some of our reads might derive from novel miRNAs not annotated in any database and used miRDeep2 (40) to investigate this possibility. The prediction performed by this tool is based on the putative miRNA primary structure and how reads are aligning to the precursor based on their biogenesis. With this assumption, reads coming from a putative novel miRNA will fall into three main categories: the mature sequence, the hairpin loop and the star sequence (22nt sequence resulting from the removal of the loop that is not loaded into Ago and degraded). If the combination of a possible hairpin precursor and mapping of the sequencing reads is not following this expected pattern, those reads are discarded. This resulted in the prediction of 163 putative novel miRNAs sequences (read count ≥1) that for convenience were labeled as miR-n-followed by a progressive number as identifier (**Figure 1a** and **Supplementary File S1**).

Next, we sought to select and validate some of these predicted miRNAs. To this end, we first chose those showing a higher consistency in detection among biological replicates (i.e. at least 2 out of 3 samples from the same cell type) reducing our initial list of 163 to 22 candidates. Next, we rank-ordered this refined cohort of putative novel miRNAs based on their average expression across cell populations selecting the top 8 for validation by Northern blot with radioactive probes (**Supplementary Table 1**). Among these, we confirmed the expression of 5 showing a size in the range of 90-150 nt (**Supplementary Figure S1a**; note that in some lanes two miR-n are probed with identical sequence but derived from different loci) which is inconsistent with the known size of either mature or pre-miRNA (20-25 nt and ∼60 nt, respectively) and more in line with that of other small RNAs including t-, sn- or sno-RNAs. Although not excluding the possibility that other novel miRNAs might be present in our list, this exclusion of 8 out 8, top-ranking putative novel miRNAs made us conclude that our catalog of mouse cortical miRNAs is virtually complete.

As a next step, we validated the robustness of our datasets following a two-step approach. First, we normalized read numbers using DESeq2 (median-ratio normalization) (41) to account for differences in sequencing depth. Upon normalization, principal component analysis (PCA) showed a clear separation of the three cell types, which distributed according to lineage differentiation (PP→DP→N) for the component displaying the highest variance (PC1) (**Figure 1b**). Second, we selected 6 miRNAs known to play key roles in neurogenesis and compared their normalized expression measured by deep sequencing with their tissue distribution assessed by *in situ* hybridization (ISH) data from Eurexpress (46). We observed a nearly perfect overlap between our sequencing and ISH data in all cases, regardless of whether the miRNA was uniformly expressed throughout the cortex (miR-9-5p and miR-17-5p), enriched in either progenitors (miR-92b-3p, miR-92a-3p and let-7b-5p) or neurons (miR-124-3p) (**Figure 1c** and **Supplementary Figure S1b**).

Taken together, our results provide evidence for an overall complete catalog of miRNA expression in cortical cell types during mouse development, more than doubling the previously known list of 292 cortical miRNA precursors (30) by detecting 421 additional ones and for a total of 703 transcripts.

### Differentially expressed miRNAs

The fine resolution of our system gave us the opportunity to assess differential miRNA expression at single population level during lineage commitment (**Supplementary File S2**). Therefore, by comparing the PP-DP and DP-N transitions, we identified miRNAs that were up- or down-regulated by >1.5-fold (i.e. log2 fold change ≥ 0.58 or ≤ −0.58, respectively, FDR <5%) in one cell type compared to its parental population. As observed previously for linear and circular transcripts (31,35), only a small fraction of miRNAs showed a significant change between PP-DP (7%) and DP-N (17%) while the majority of those up- or down-regulated between PP-DP continued to follow the same trend of up- or down-regulation, respectively, between DP-N (**Figure 2**).

**Figure 2.**
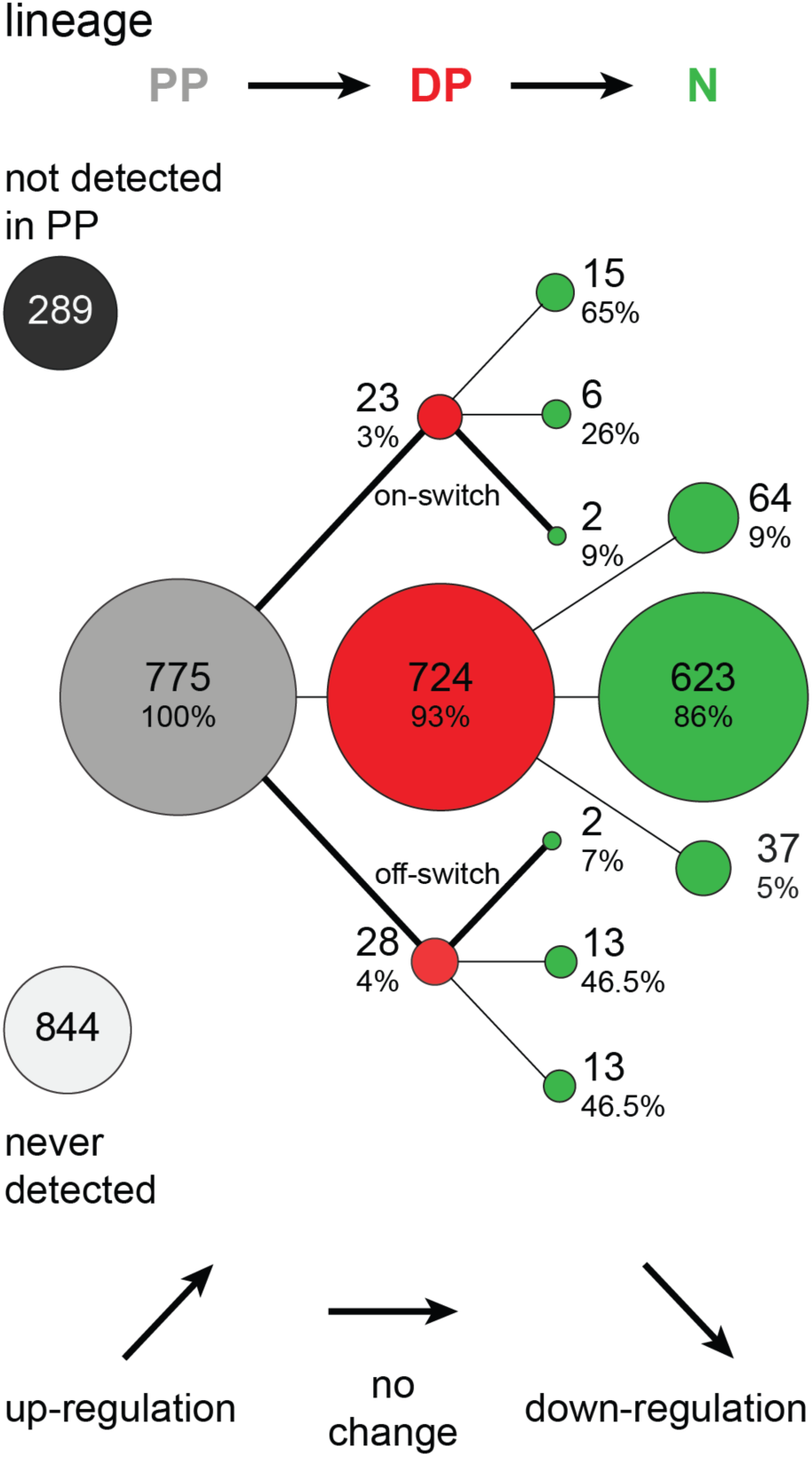
Differential expression analysis. Representation of differentially expressed miRNAs in the three cell types (PP: grey; DP: red; N: green). Numbers indicate the number of miRNA in each group and percentages are calculated over the parental population. miRNAs not detected in PP or never detected in any cell type are also reported (top left and bottom left, respectively). Oblique lines represent a > 50% change (log_2_ fold change ≥ 0.58 or ≤ −0.58) and FDR <5%, whereas horizontal lines a < 50% change or an FDR >5%. Bold lines are depicting on- and off- switch patterns.

Furthermore, by analyzing coding and long non-coding transcripts, our group previously concluded that a transient up- or down-regulation specifically in DP compared to both their PP progenitors and N progeny (on- and off-switch transcripts, respectively) represented a hallmark of functional commitment to the neurogenic lineage (31,36). Intriguingly, the subset of miRNAs displaying this on-/off-switch pattern of expression was strongly underrepresented, accounting for only 0.5% of the total and suggestive of a highly specific expression pattern. In fact, we only found 2 on-switch (let-7b-5p and miR-135a-2-3p) and 2 off-switch (miR-486a-5p and miR-486b-5p) miRNAs (**Figure 2** and **Figure 3**). Supporting our conclusion that on-/off-switch transcripts are functionally involved in neurogenic commitment, both let-7b and miR-135a-2 are well known to be key regulators of neurogenesis (17,47). In contrast, while it has been shown that miR-486a and miR-486b promote myoblast differentiation (48) and are involved in regulatory pathways of ectodermal-derived tissues (49,50), no neurogenesis-related function has ever been reported for these two miRNAs to date.

**Figure 3.**
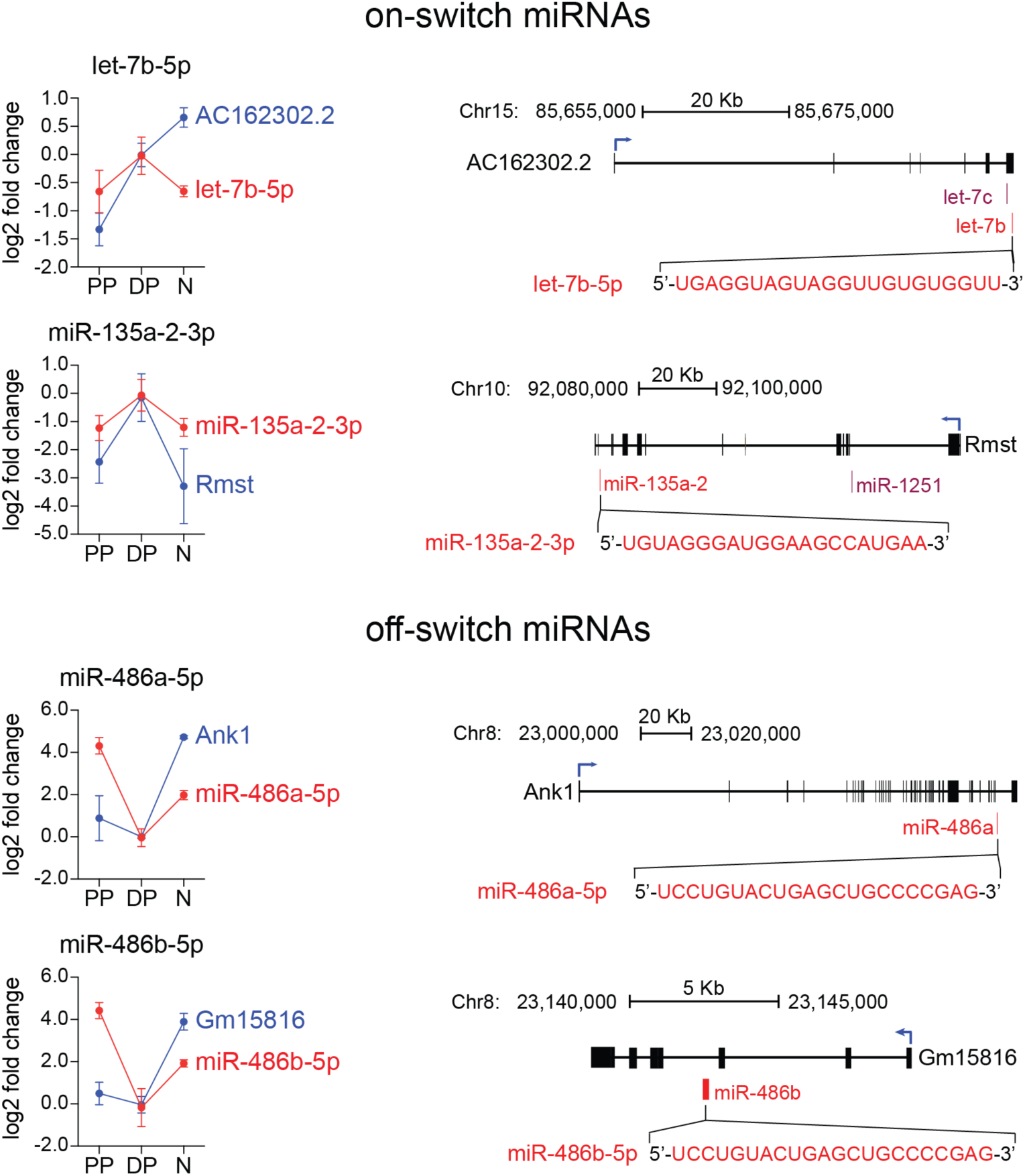
Genes hosting switch miRNAs are depicted (black): blue arrows represent the direction of transcription, whereas black boxes and lines constitute exons and introns, respectively. Position and mature sequence of switch miRNAs are indicated in red. Expression patterns of miRNAs and host genes are reported (graphs) on the left.

With regard to their genomic location, we observed that all 4 switch miRNAs were intragenic. In particular, let-7b-5p and miR-135a-2-3p were processed, respectively, from lncRNAs AC162302.2 and Rmst (which mediates Sox2-dependent progenitors proliferation) (51). Similarly, miR-486a-5p and miR-486b-5p were processed from Ankirin1 (Ank1) and the predicted gene Gm15816, respectively. Interestingly, miR-486a-5p and miR-486b-5p shared the same mature sequence, despite originating from different pre-miRNAs transcribed from opposite strands of the same genomic locus. Unsurprisingly, when analyzing the expression pattern of the host genes of switch miRNAs (data retrieved from (31)), we found a high degree of overlap in their differential expression within different cell populations (**Figure 3**). This was consistent with previous observations of our group on long non-coding and circular RNAs (cite-cite) in which intragenic, switch genes were found to be regulated together in a similar fashion at the level of a common “switch locus”.

### The off-switch miR-486a/b-5p is a novel regulator of neurogenesis

The absence of any known function for miR-486a/b-5p in neural stem cells or brain development, together with their intriguing switch expression pattern, drove us to investigate their potential functional role during corticogenesis and, by this, also attempting to validate the power of our miRnome atlas of cortical cell types. After validating the expression of miR486a/b-5p by qRT-PCR on FAC-sorted PP, DP and N of the E14.5 cortex (**Supplementary Figure 2a**), we used locked nucleic acids (LNA) to inhibit their activity and, hence, address their functional role. First, we confirmed the silencing efficacy of LNA-486 by luciferase assay on two validated targets of miR-486a/b-5p (Foxo1 and Pten) (52,53) (see Materials and Methods and **Supplementary Figure S2b** and **c**). Then, to investigate the effect of miR-486a/b-5p inhibition on cortical progenitors, we *in utero* electroporated E13.5 mouse embryos with LNA-486 or LNA-control together with an RFP-reporter plasmid. Brains were collected 48h later and distribution of electroporated cells (identified as RFP+) was assessed as a readout of neurogenesis and neuronal migration.

LNA-486 significantly altered cell distribution across all cortical layers. Particularly affected were the subventricular zone and the cortical plate that showed a 1.6 fold increase (from 12±1 to 20±2%; p<0.01), and a comparable decrease (from 20±3 to 12±2%; p<0.01), in RFP+ cells after delivery of LNA-486 relative to brains electroporated with control LNAs, respectively (**Figure 4a**). By using Tbr2 as a marker to identify basal from apically progenitors, we observed a significant increase in both cell types at the expense of neurons. In particular, apical progenitors increased by 1.2 fold (from 14±0.2 to 18±2%; p<0.05), whereas basal increased by 1.3 fold (from 21±2 to 28±4%; p<0.05). This was paralleled by a comparable, 1.2 fold decrease in neurons found in the neuronal layers (from 66±2 to 54±2%; p<0.001) (**Figure 4b**). Notably, no major effect was found neither at the level of cell survival nor migration of newborn neurons as assessed by activated-caspase immunoreactivity or upon 24h birthdating with BrdU (data not shown) and hinting at a cell fate-specific effect upon inhibition of miR-486a/b-5p activity.

**Figure 4.**
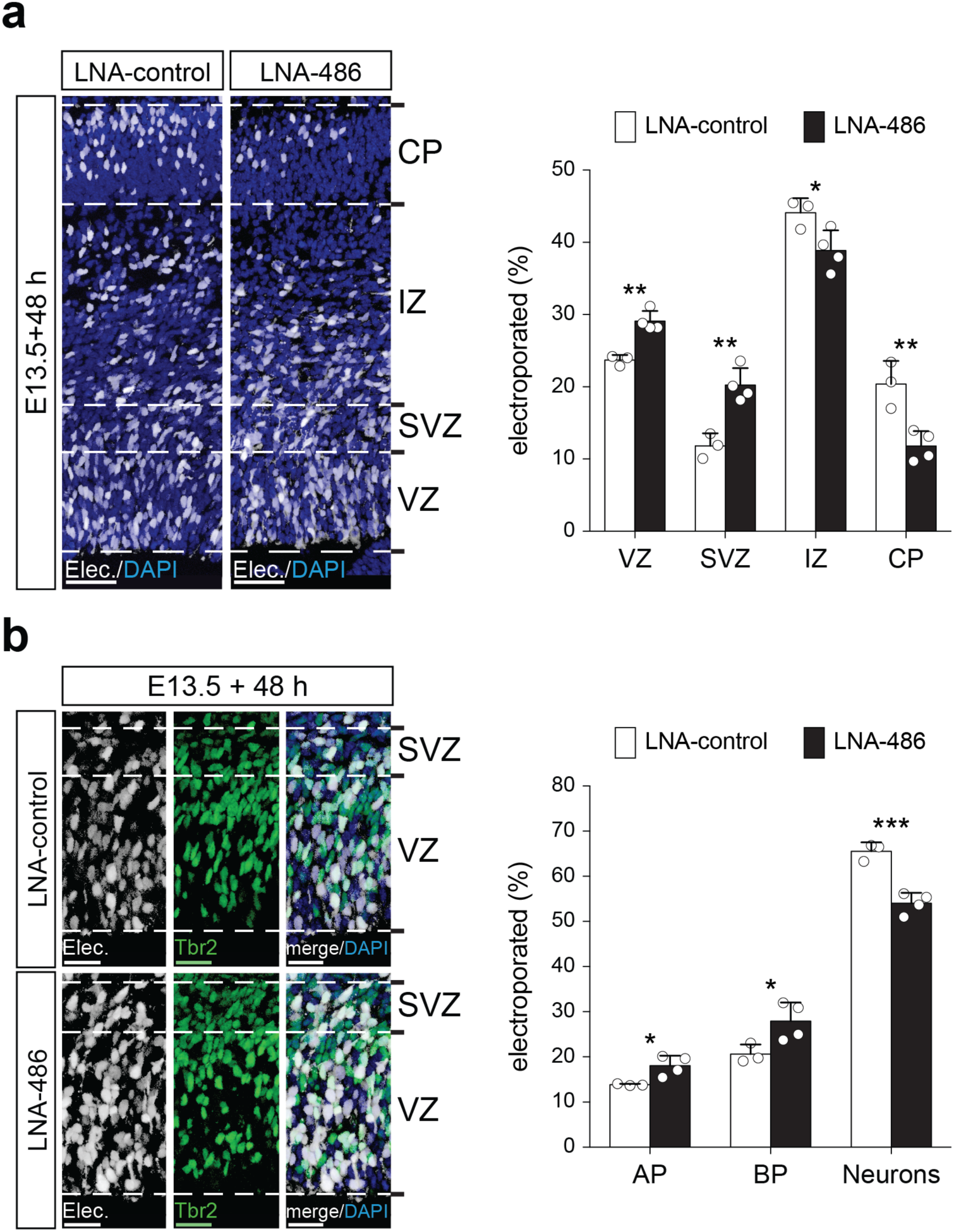
Manipulation of miR-486a/b-5p. (**a**,**b**) Coronal sections of electroporated lateral cortices stained for RFP (electroporated cells, white), DAPI (all nuclei, blue) (**a**) and Tbr2 (BP, green) (**b**). Histograms represent quantifications of cells distribution 48 hours after electroporation of LNA-control (white) or LNA-486 (black) (histograms). VZ: ventricular zone; SVZ: sub-ventricular zone; IZ: intermediate zone; CP: cortical plate. AP: apical progenitor (Tbr2-cells in VZ); BP: basal progenitor (Tbr2+ cells in VZ and SVZ). Error bars = s.d. N ≥ 3. Individual dots represent biological replicates. *p<0.05; **p<0.01; ***p<0.001. Scale bar = 25 µm.

Taken together, characterization of the miRNome of progenitor cell types and neurons revealed a powerful tool to identify new miRNAs involved in cortical development allowing us to describe for the first time the functional effects of interfering with the activity of miR-486a/b.

## DISCUSSION

Here we provided a complete catalog of miRNAs expression in neural progenitors and newborn neurons during cortical development and validated our resource by identifying a new player in neural stem cell fate specification: the switch miR-486a/b-5p.

Since the discovery of miRNAs as critical regulators of translation (7), many groups attempted to obtain atlases of their expression during brain development. The use of microarrays offered a first approach toward this goal (26-29) but was limited by a previous knowledge about the sequence of such miRNAs. Next-generation sequencing overcame this limitation and significantly increased the number of known miRNAs (30). However, previous studies remained limited either to the use of cell cultures or to analyses of whole brain lysates due to a lack of systems to discriminate between different cellular subtypes coexisting in time and space during corticogenesis. Even with the advent of single-cell sequencing, the study of small RNAs remains hindered by two major technical limitations that a) drop-seq is currently applicable only to poly(A)-RNAs and b) library preps with <1,000 cells display extremely poor coverage (25).

Here we exploited the Btg2::RFP/Tubb3::GFP mouse line as a well-established tool used by our group in previous studies to characterize the molecular signature of neurogenic commitment (34-37). By doing so, our group identified switch transcripts belonging to several classes of RNAs and including coding, long non-coding and circular RNAs and in most cases showing their functional roles in brain development (31,35,36). Continuing this line of research, here not only we provided the field with a validated and overall complete catalogue of cortical miRNAs at single-population level but also identified in miR-486a/b-5p a novel regulator of neurogenesis. We hope that future studies will be able to dissect the molecular mechanisms underlying this novel cortical, switch miRNA and that the field in general will profit from this novel resource.

## ACCESSION NUMBERS

Sequencing data generated during the current study are available at GEO repository (GSE142253)

## ACKNOWLEDGMENTS

We thank the MPI-CBG and CRTD facilities for maintenance of the mouse lines, sequencing and FACS.

## AUTHORS CONTRIBUTION

FC, MD and DC conceived the project; MD carried out the bioinformatic analyses (supported by ML and AD) and DC performed experiments with the help of and SM, LHAA and BCT. SK and GS performed the Northern Blot. FC, MD and DC wrote the manuscript. All authors approved the manuscript.

