## Supplementary Data for "microRNA profiling of mouse cortical progenitors and neurons reveals miR-486-5p as a novel regulator of neurogenesis"

#### SUPPLEMENTARY MATERIALS AND METHODS

##### Northern blot

30 µg of total RNA extracted from E14.5 mouse cortices were separated using denaturing urea 15 % PAGE gel (Mini-PROTEAN system; Bio-Rad) in 1x TBE and blotted onto a GeneScreen Plus nylon membrane (PerkinElmer) in pre-cooled 0.5x TBE. Radioactively labeled Decade marker (Ambion) was used as molecular marker. RNAs were cross-linked to the membrane by UV irradiation (1,200 mJ), followed by baking for 30 min at 80 °C. The membrane was pre-incubated in hybridization buffer (5xSSC, 20 mM $Na_2HPO4$ (pH=7.2), 7 % SDS, 2 x Denhardt's solution, 40 µg/mL salmon sperm DNA) for 2 h at 50 °C in constant rotation, followed by incubation overnight at 50 °C in hybridization buffer containing the denatured [32P] labeled DNA probes against the predicted novel mature miRNA sequences. Probes against miR-9-5p and miR-124-3p, were used as positive controls. The membrane was washed twice for 10 min and twice for 30 min at 50 °C with non-stringent wash solution (3 x SSC, 25 mM $NaH_2PO4$ (pH=7.5), 5 % SDS, 10 x Denhardt's solution) and once for 5 min at 50 °C with stringent wash solution (1 x SSC, 1 % SDS). Signals were detected by autoradiography using the Cyclone Plus Phosphor Imager (PerkinElmer). The membrane was stripped (0.1 % SDS, 5 mM Na-EDTA, preheated to 95 °C) for 1 h and re-used several times to detect additional miRNAs.

##### Immunohistochemistry

Cryosections were permeabilized (0.5 % Triton X-100 in PBS) for 20 min, quenched (0.1 M glycin in PBS) for 30 min and blocked (10 % donkey serum, 0.3 % Triton X-100 in PBS) for 30 min at RT. All primary antibodies were incubated (3 % donkey serum, 0.3 % Triton X-100 in PBS) overnight at 4˚C, followed by washing and incubation with secondary antibodies for 2 h at RT (See **Supplementary Table 2** for a list of used antibodies). For Tbr2 staining, antigen-retrieval was performed in 0.01 M Citrate Buffer (pH=6.0) for 1 h at 70˚C.

##### *In situ* hybridization

*In situ* hybridization was performed as previously described (54), using digoxigenin (DIG) – labeled LNA probes purchased from Exiqon (Sequences listed in **Supplementary Table 1**). Cryosections were fixed (4 % PFA) for 10 min and acetylated (100 mM triethanolamine, 0.25 % acetic anhydride) for 15 min constantly rocking at RT. Pre-hybridization was carried out in hybridization buffer (HB) (50 % formamide, 5X SSC, 0.1 mg/ml Heparin, 1X Denhardt solution, 0.1 % CHAPS, 0.2 mg/ml yeast tRNA, 0.01 M EDTA, 0.4 % Tween-20 in DEPC-H2O) for 1 h at 53˚C. Hybridization was performed in HB containing 50 nM LNA probes (previously denatured at 75˚C for 5 min) overnight at 53˚C. Sections were washed first in washing buffer (50 % formamide, 2X SSC, 0.1 % Tween-20) for 90 min at 53˚C, then in 2X SSC for 1 h at 53˚C and eventually in maleic acid buffer (MABT) (100 mM maleic acid, 150 mM NaCl, 0.25 % Tween-20, pH=7.8) for 30 min at RT. Next, blocking solution (MABT, 20 % goat serum, 2 % Boehringer Blocking Reagent) was applied for 30 min at RT, followed by incubation with anti-DIG-AP antibody (1:2000 in blocking solution, Roche) overnight at 4˚C. Washings were performed in MABT for 90 min and NTMT (100 mM Tris-HCl pH=9.5, 100 mM NaCl, 50 mM $MgCl_2$, 0.1 % Tween-20) for 1 h at RT. Labeling was developed in BM purple (Roche) overnight at 37˚C. Sections were imaged using an automated microscope (ApoTome; Carl Zeiss), pictures digitally assembled using Zen software (Carl Zeiss) and composites analyzed using Photoshop CS6 (Adobe).

**Cloning and luciferase assay**

psiCHECK-2 double luciferase vector containing human PTEN 3'-UTR flanked by XhoI and NotI restriction sites was purchased from Addgene (cat #50936). Human Pten 3'-UTR was replaced by parts of the 3'-UTRs of Foxo1 and Pten, which were PCR-amplified from mouse genomic DNA (kindly provided by Sara Zocher, Kempermann G. lab) and inserted into the psiCHECK-2 vector downstream of Renilla Luciferase. For Foxo1, the 3'UTR region between nucleotides 12 – 1281 was amplified, while for Pten, the region between nucleotides 2558 – 3865 was selected (primers pairs are listed in **Supplementary Table 1**). Mutants of both constructs carrying a 3-nt-mutation in miR-486-5p binding site were generated in two steps. First, each cloned 3'UTR was amplified in two different (but overlapping) fragments carrying the mutation. Then, the fragments were pooled and re-amplified using the cloning primer pairs as above. The generation of a vector expressing pre-miR-486-5p under control of a U6 promoter and a nuclear-localized red fluorescent protein (RFPnls) under control of a SV40 promoter was done as follows. The whole RFPnls cassette (SV40 promoter, RFP coding sequence, nls and poly-adenylation signal) was excised from pDSV-RFPnls plasmid (52) using SspI restriction enzyme (NEB) and cloned into the pSilencer™2.1-U6-Neo vector (Thermo Fisher) (replacing the

Neomycin cassette). pre-miR-486a (miRBase accession number: MI0003493) plus 50 bp flanking each side were PCR-amplified from mouse genomic DNA (kindly provided by Sara Zocher, Gerd Kemperman's Lab) and cloned downstream of the U6 promoter using BamHI and HindIII restriction sites. Primers for miR-486a are listed in **Supplementary Table 1**. All sequences (and introduced mutations) were confirmed by Sanger sequencing. Luciferase assays were performed using Neuro2A (N2A) cells maintained in DMEM (Gibco) supplemented with 10 % FBS at 37˚C and 5 % $CO^2$. 7 x $10^5$ cells/well were seeded in 24-well plates and co-transfected with 215 ng of psiCHECK-2, 150 ng of miR-486a plasmid and 100 – 150 nM LNA, using polyethylenimine (PEI, Sigma Aldrich) (PEI:DNA ratio 3:1). 24 hours after transfection cells were washed with PBS, lysed and luciferase assay was performed using Dual Luciferase Reporter Assay System (Promega).

#### SUPPLEMENTARY TABLES AND FIGURES

**Supplementary Table 1**. List of LNA, primers and oligonucleotide sequences

| Experiment | Oligo name | Sequence (5' → 3') |
|---|---|---|
| IUE; Luciferase assay | LNA-control | TAACACGTCTATACGCCA |
| | LNA-486 | TCGGGGCAGCTCAGTACAG |
| *In situ* hybridization | LNA-control | DIG-TCACTGCATACGACGATTCT |
| | LNA-486 | AACCACACAACCTACTACCTCA-DIG |
| Northern Blot | mmu-miR-n-12 | TGGGGGCTCGTTCGGGAT |
| | mmu-miR-n-68 | TGGGGGCTCGTTCGGGAT |
| | mmu-miR-n-37 | TCTGCTCTGAGGTCGAGG |
| | mmu-miR-n-19 | GCTCTACCACTGAGCTACATCCCC |
| | mmu-miR-n-29 | TATTGTCTGAAATTTCTAT |
| | mmu-miR-n-5 | TATTGTCTGAAATTTCTAT |
| | mmu-miR-n-47 | TGGAGGCCCCAGCGAGAT |
| | mmu-miR-n-28 | CATGCAGAGTGAGAGAGT |
| | mmu-miR-9-5p | TCATACAGCCTAGATAACCAAAGA |
| | mmu-miR-124-3p | GGCATTCACCGCGTGCCTTA |
| UTR Cloning (red depicts RE sites) | Foxo1_XhoI_fwd | GGTTCTCGAGAGGCTACATTTAAAAGTCCTTC |
| | Foxo1_NotI_rev | ATTAGCGGCCGCACAAAGAACATCACCTTAG |
| | Pten_XhoI_fwd | GGTTCTCGAGCTCCCGTGTTTCTGGAATGC |
| | Pten_NotI_rev | ATTCGCGGCCGCTCATGTAACATTAAGACTCC |
| UTR mutation (red depicts mutation site) | Foxo1_mut_fw | GATTAAGTGCCAGCTTTGTTGTGGTCTTTTTCTAT TGTTTTTGTTGTTGTTTATTTTGTT |
| | Foxo1_mut_rev | AACAAAATAAACAACAACAAAAACAATAGAAAAA GACCACAACAAAGCTGGCACTTAATC |
| | Pten_mut_fw | GATACACAAATATGACGTGTTGTGGATAATGCCT CATACCAATCAGATGTCCATTTGTTA |
| | Pten_mut_rev | TAACAAATGGACATCTGATTGGTATGAGGCATTA TCCACAACACGTCATATTTGTGTATC |
| miR cloning (red depicts RE sites) | pre-miR-486_BamHI_fwd | ATCGGGATCCTACGCAACGAAGATCTTCAGCAG |
| | pre-miR-486_HindIII_rev | TGCCAAGCTTTTCAAAAAGAAGGGGCAATAACCC AGTTAG |

**Supplementary Table 2**. List of antibodies used for immunohistochemistry

| Antigen | Supplier | Cat n.° | Species | Dilution |
|---|---|---|---|---|
| RFP | Chromotek | 5F8 | Rat | 1:400 |
| RFP | Rockland | 600-401-379 | Rabbit | 1:2000 |
| Tbr2 | Abcam | ab183991 | Rabbit | 1:500 |
| Secondary Antibodies | Jackson | 712-165 -150<br>711-165-152<br>711-545 -152 | Donkey | 1:500 |

### Suppl. Figure 1 Dori M. & Cavalli D. *et al.*,

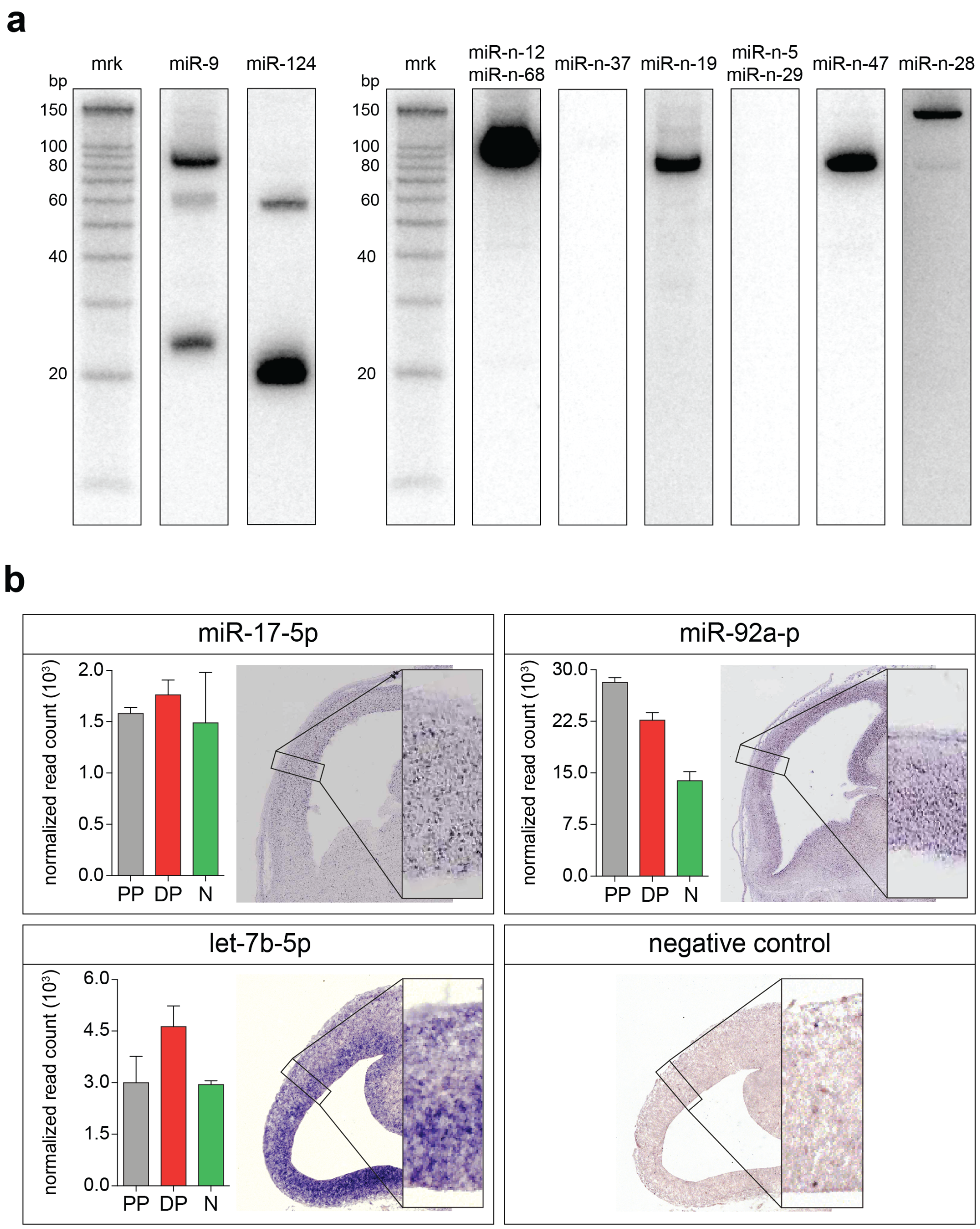


**Supplementary Figure 1.** Validation of novel and known miRNAs. **a)** Northern blots performed with 32P-labeled DNA probes on RNA from E14.5 cortices. Radioactive markers (mrk) were use to determine fragment size and mature miRNA names are reported in boxes (top). Double names indicate identical mature miRNA sequence. miR-9 and miR-124 were used as positive controls. The 80 nt band in miR-9 is a carryover from hybridization with miR-n-19 probe (despite stripping). All blots were performed in biological triplicates. **b.** Sagittal sections downloaded from Eurexpress (miR-17-5p, miR-92a-3p) or coronal sections hybridized in house (let-7b-5p, Negative control) of E14.5 cortices. Magnifications of the lateral cortex are shown (bottom-right) to appreciate the extent of the overlap with miRNA expression data measured by deep sequencing (histograms). Error bars = s.d. N = 3.

### Suppl. Figure 2 Dori M. & Cavalli D. et al.,

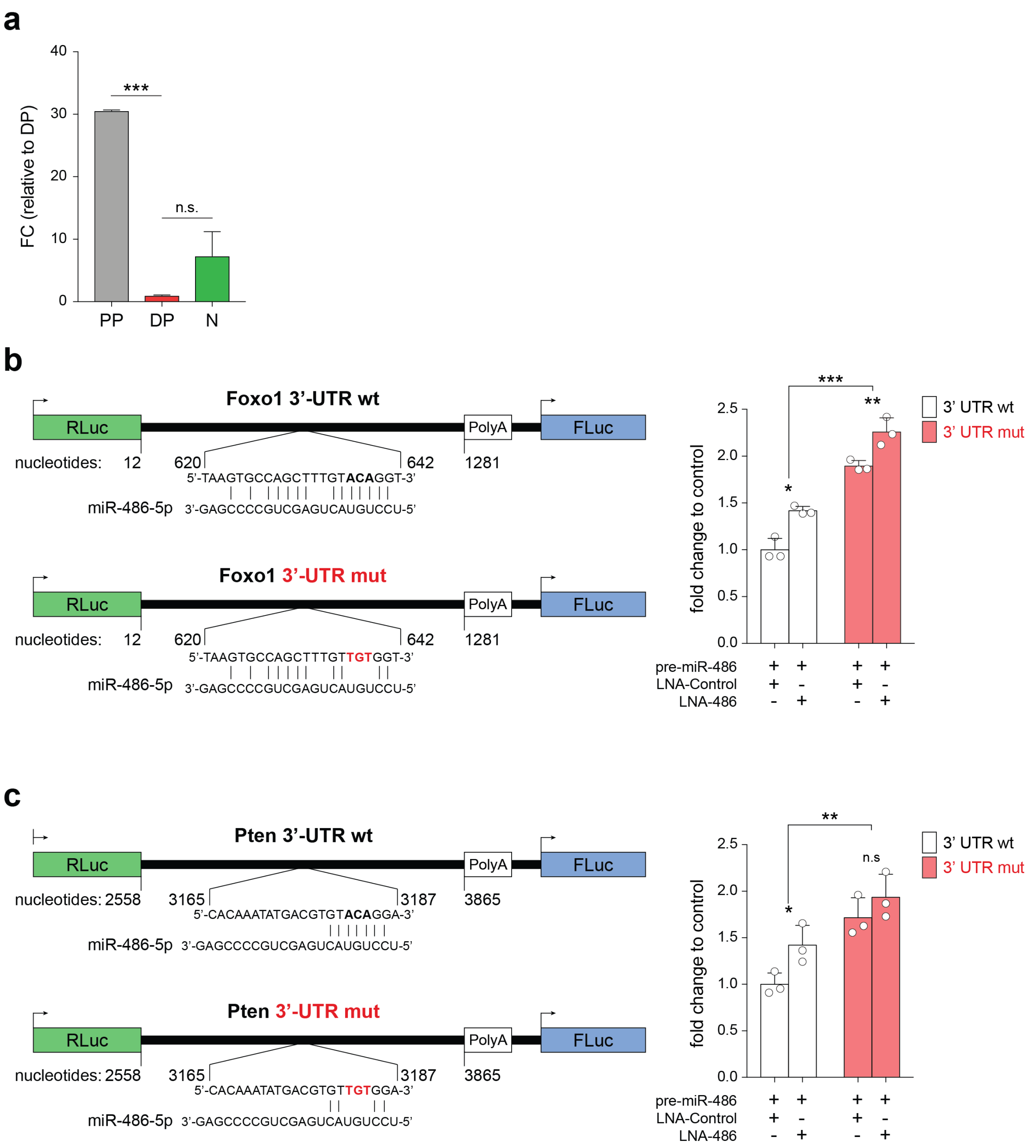


**Supplementary Figure 2.** (a) miR486a/b expression was validated through qPCR on RNA from sorted cells obtained from E14.5 cortices. Validation of miR-486a/b-5p knockdown by luciferase assay. psiCHECK-2 vectors containing mouse Foxo1 (b) or Pten (c) 3'-UTR are reported (left). Nucleotide numbers refer to the portion of the 3'-UTR used for cloning. Binding sites of miR-486a/b are reported (drop down) and mutated nucleotides are marked in red. Luminescence quantifications (histograms) for wild type (white) and mutated (red) constructs co-transfected with plasmid expressing the pre-miR-486 and either LNA-Control or LNA-486 (x-axis) are expressed as fold change (y-axis) relative to control (3'-UTR wt + LNA-Control). Error bars = s.d. N ≥ 2. Individual dots represent biological replicates. *p<0.05 ; **p<0.01; ***p<0.001.
